# Neural markers of visual working memory during an upper extremity movement-coupled task: a mobile brain-body imaging (MOBI) study

**DOI:** 10.64898/2026.01.04.697538

**Authors:** Jeehyun Kim, Edward G. Freedman, John J. Foxe

**Author notes:** Corresponding & Co-Senior Authors &.

## Abstract

Optimal goal-directed behavior in everyday life involves the appropriate coupling of cognition and movement. In this study we employed a mobile brain-body imaging (MoBI) system that integrated time-synchronized high-density electroencephalography with 3-dimensional motion capture during a touchscreen visual working memory task requiring dynamic upper extremity reach responses. Using this framework we examined how increasing memory load (i.e. from 1-disc to 4-discs) influences behavior and scalp-recorded neural activity when cognitive and motor processes are coupled within a single task and whether these effects are modulated by handedness. Increasing memory load reliably degraded working memory task performance and slowed motor preparation, while core movement execution measures remained largely preserved. Regarding electroencephalography data, parieto-occipital activity increased with load during both the early visual-evoked N1 and the delay period. However, load-related modulation reached a plateau at the 3-disc condition during the delay period only. Posterior-frontal engagement also strengthened as load increased, consistent with demands on attentional control, memory maintenance, and sensorimotor planning. Brain-behavior associations were most robust at a moderate level (2-disc), where contralateral delay activity correlated with both cognitive and motor performance measures. Finally, although handedness exerted selective behavioral effects, it did not modulate load-related parieto-occipital neural activity. Together, these findings demonstrate that a multimodal MoBI approach captures integrated cognitive-motor dynamics during working memory-guided actions, providing a more comprehensive characterization of behavior and underlying neural signatures than either modality alone. This framework offers a translational approach for studying cognitive-motor interactions and their potential disruption in neurological conditions such as Parkinson’s disease.

## 1. INTRODUCTION

Visual working memory (WM) enables temporary maintenance of object features and locations to guide upcoming goal-directed behavior, and its performance predicts outcomes across diverse cognitive domains, underscoring its role as a core construct essential to a broad range of behaviors [1, 2]. Behavioral studies of visual WM consistently show robust load effects, with capacity plateauing at roughly three to four items [3–5].

Most prior evidence comes from paradigms using simplified motor outputs, such as button presses, which do not capture the naturalistic dynamics of movements like reaching. This raises the question of how increasing WM load shapes the perceptual, mnemonic, and preparatory processes that precede more naturalistic behavioral outputs when memory and action are coupled in a single task.

Electroencephalography (EEG)-based event-related potentials (ERPs) provide millisecond-level access to neural activity driving actions. Over posterior parieto-occipital cortical regions, the early visual-evoked N1 component (∼ 150–200 msec post-encoding array onset) indexes selective sensory processing [6–8], while the sustained delay period negativity (∼ 300–900 msec post-encoding array onset), such as contralateral delay activity (CDA), indexes maintenance, scaling with the number of items until capacity is reached [3, 9]. Dipole modeling and intracranial recordings localize the N1 to lateral occipital cortical regions [7, 8, 10–12], whereas the CDA is thought to reflect activity in posterior parietal cortex (PPC) region [3, 9, 13]. Together, these signatures serve as canonical neural markers of perceptual selection and WM maintenance that can be examined prior to task-related performance.

Electrophysiological and neuroimaging studies have shown that parieto-occipital regions support early visual processing [14] and maintenance that plateau at capacity [3, 13], while frontal regions, such as the prefrontal cortex (PFC), become critical for consolidation and executive control, especially at higher demands [15–19]. Beyond sensory processing and maintenance, activity within the PPC contributes to the encoding of intentions and, along with the frontal regions, mediates the transformation of sensory input into prospective motor plans, supporting sensorimotor integration and movement preparation [20–25]. Further, delay period activity in the PFC and PPC likely reflects a convergence of attention, WM maintenance, and movement intention [26], underscoring the broader functional role of the frontoparietal network. Considering that these functions draw on shared resources, we hypothesize that increasing memory load consumes more of this shared capacity, thereby impairing motor planning and, in turn, motor execution. To better capture such dynamics, we employ a mobile brain-body imaging (MoBI) system, which records time-synchronized neural and kinematic signals during a visual WM task that couples dynamic arm movements as behavioral outputs [27]. This approach allows us to track WM load effects across distinct task phases and delineate where demand imposes costs in the sensorimotor pipeline.

Handedness introduces an additional axis of interest. Using the non-dominant hand may impose additional planning and control complexity before movement onset, particularly under high WM load. In our action-coupled visual WM task, these asymmetries could manifest as changes in motor preparation and kinematic control. Testing whether canonical posterior signals, such as N1 and CDA, remain invariant across hands, especially when targets and response hands fall on the same versus opposite sides, will clarify whether these neural markers primarily index mnemonic and attentional processes or are modulated by handedness-specific motor control.

To address these questions, we use high-density EEG during a touchscreen visual WM task that requires memorizing one to four items and subsequently reaching to the remembered location using either the dominant or non-dominant hand. By embedding this design within a MoBI system, we can capture time-synchronized neural and kinematic signatures in a naturalistic, action-coupled setting. This approach not only simulates everyday cognition-action coupling more closely than classic button-press paradigms but also enables direct examination of load effects prior to behavioral performance. Guided by prior work, we hypothesize that: (1) increasing the load will degrade memory performance, prolong motor preparation, and worsen movement performance; (2) parieto-occipital contralateral ERP activities will scale with load during both the early visual-evoked period and subsequent delay period, plateauing at capacity only during the delay period, with complementary frontal and central positivity consistent with frontoparietal engagement; (3) posterior load-related activity will correlate with task-related behavioral performances; and (4) handedness will exert selective effects on behavioral performances and posterior load-related activity. By integrating continuous behavioral measures with lateralized pre-movement ERPs, this study aims to elucidate how visual WM load affects the sensorimotor pipeline, whether posterior neural signals extend to action-coupled, handedness-engaged contexts, and the load level at which WM-related signals best predict behavior.

## 2. MATERIALS & METHODS

### 2.1. Participants

Thirty-two individuals with normal or corrected-to-normal vision were recruited from the University of Rochester and the surrounding community. Six participants were excluded from analysis – 2 due to loss of follow-up and 4 due to data quality and/or technical issues affecting data acquisition. The final analysis group included 26 participants (mean ± standard deviation (SD): age = 24.2 ± 5.4 years; education = 16.2 ± 3.1 years; 19 female, 7 male; 24 right-hand dominant, 2 left-hand dominant). No participants reported any history of psychiatric, neurological, or musculoskeletal disorders, recent head injuries, or history of substance abuse. We conducted cognitive screening with the Montreal Cognitive Assessment (MoCA) [28], and all participants scored ≥20, a threshold associated with optimal diagnostic accuracy for cognitive impairment [29]. Participants also completed the Wechsler Adult Intelligence Scale (WAIS)-IV Digit Span [30], Edinburgh Handedness Inventory [31], and an experimental data acquisition session. The Institutional Review Board of the University of Rochester approved the experimental procedures, and all participants provided their written informed consent. All procedures were compliant with the principles laid out in the Declaration of Helsinki for the responsible conduct of research, with the exception that this study was not pre-registered. Participants were paid $18/hour for time spent in the lab.

### 2.2. MoBI System Setup

All behavioral, neural, and kinematic data were time-synchronized using Lab Streaming Layer (LSL) software (Swartz Center for Computational Neuroscience, University of California, San Diego, CA, USA; available at: https://github.com/sccn/labstreaminglayer) that integrated touchscreen-based behavioral response, high-density EEG, and 3-dimensional (3D) motion capture data. Past studies have demonstrated the reliable use of the MoBI technique for examining neural activity associated with a range of cognitive and motor functioning, and its long-term test-retest reliability has been shown [27, 32–37].

Participants were seated in front of a platform consisting of a touchscreen monitor (Model 22TS5M, Beetronics; 24-inch, resolution 1920 x 1080 pixels) and a central weight-sensitive homebase (Arduino microcontroller with HX711 converter and calibrated load cell), which ensured uniform hand placement prior to trial initiation. The distance between the central homebase and the touchscreen monitor was kept constant across all participants (∼ 117 cm). We ensured that participants could view the entire screen with their eye level aligned with the center of the monitor and measured arm lengths (i.e., from acromion to the tip of index finger; mean ± SD: 68.9 ± 3.5 cm (right), 68.8 ± 3.8 cm (left)).

### 2.3. Behavioral Task Paradigm

Participants completed the behavioral task shown in **Figure 1**. We designed the task to probe cognitive-motor integration using both memory trials (i.e., trials with varying memory loads and motor responses) and motor-only trials (i.e., trials with only motor responses with no memory demands). The behavioral task was controlled by custom software written in Presentation (Neurobehavioral Systems Inc., Berkeley, CA, USA), which also generated behavioral event markers relating to stimulus triggers and behavioral responses. Participants were seated ∼ 50 cm from the touchscreen (i.e., eye-to-screen distance) and instructed to place their response-hand (i.e., the side designated for that session) on the central homebase before each trial. Each trial initiated only when the participant’s hand rested on the homebase with pressure > 50 g (mean of 10 samples). For memory trials, participants were instructed to memorize the orientation of the open-mouthed discs of varying memory loads at fixed eccentricity in the cued hemifield (i.e., target hemifield; left or right), while ignoring the same number of disc stimuli in the uncued hemifield (i.e., distractors). We presented equal numbers of stimuli on both sides to control for sensory input. Given the visual WM capacity of three to four items [3–5], set sizes varied from 1 to 4 discs per cued hemifield. Participants were instructed to fixate on the central crosshair and covertly attend to the cued hemifield when the disc stimuli appeared (i.e., encoding array; 150 msec). After encoding array offset, participants maintained central fixation during the retention interval (1,200 – 1,500 msec) until a bold target ring appeared at the location of one of the previously presented discs in the cued hemifield (i.e., recall array), signaling to respond. Using their response-hand, participants tapped with their index finger along the outline of the bold ring at the location where they recalled the center of the target disc’s opening. To minimize visual overlap, we separated disc stimuli within each hemifield by at least 180 pixels (center-to-center). **Figure 1** includes further details of the task paradigm. For motor-only trials, participants were instructed to simply tap the visible center of the opening of a single target disc, serving as a baseline measure of motor precision. We presented motor-only trials after memory trials to minimize confusion about task instructions. The task was self-paced, with each trial starting once participants rested their response-hand on the central homebase. Each participant started the first block with their dominant hand and alternated response-hands across blocks, completing up to 3 left-hand and 3 right-hand blocks. Each block lasted ∼ 10 min, including 70 memory trials (1–4 discs, pseudorandomized) and 20 motor-only trials. We visually inspected gaze position using a synchronized wearable eye-tracking system (Pupil Core, Pupil Labs GmbH, Berlin, Germany) to ensure central fixation when needed and provided feedback when necessary. We instructed participants that both speed and accuracy were equally important for this task. In between blocks, we offered a short break and reminded them as to which hand to use for the following block. Participants completed as many supervised practice runs as needed until they were comfortable with the task instructions before the recording session.

**Figure 1.**
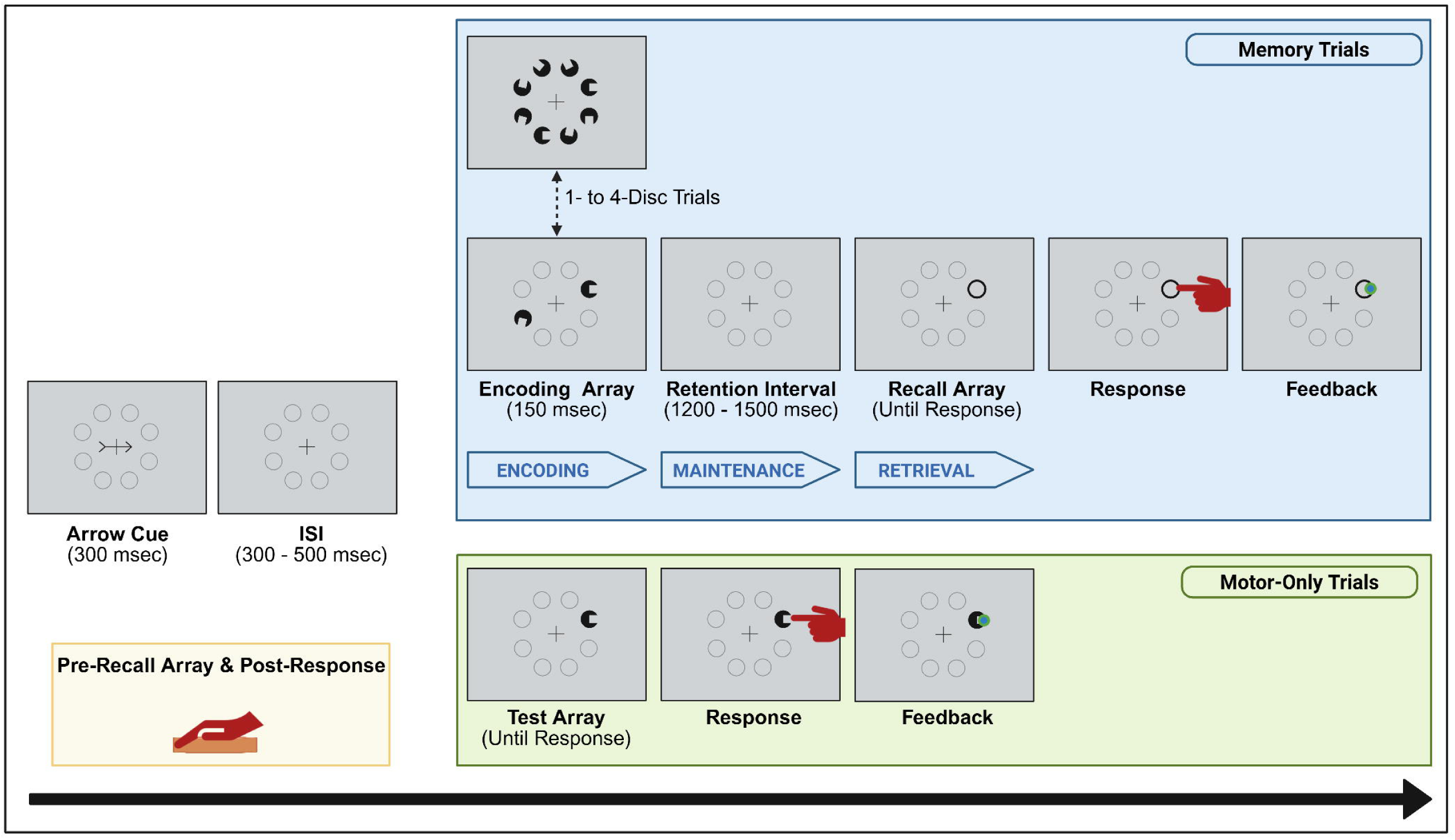
Illustration of the Behavioral Task Paradigm. In memory trials, each trial began with a central fixation cross, followed by a directional **arrow cue** (300 msec) indicating which hemifield to attend. After a jittered **interstimulus interval** (ISI; 300 – 500 msec), an **encoding array** of 1 – 4 open-mouthed discs per hemifield was presented for 150 msec. Each disc (diameter ∼3.3 cm, ∼3.2° visual angle) contained a small opening (∼1.3 cm, ∼1.2°) randomly positioned along the circumference. Stimuli were arranged symmetrically across hemifields at a fixed eccentricity of 400 pixels (∼ 10.6°) from fixation, distributed evenly along a semicircular arc. Following a 1,200 – 1,500 msec **retention interval**, during which only the central fixation cross was shown, a **recall array** appeared and remained until response or for up to 6,000 msec. Recall array was the cue for participants to initiate response-hand movement to give responses on the touchscreen monitor. One bold target ring was shown at the location of one of the previously presented disc stimuli, and participants responded by tapping the remembered center of the opening along the outline of the ring with their index finger. In motor-only trials, there were no encoding array and retention interval. Instead, after the arrow cue and ISI, a single target disc appeared and remained until response or for up to 6000 msec. Participants tapped the center of the opening with the stimulus continuously shown using their index finger, eliminating memory demands. For all trials, a small blue dot appeared at their tap location to confirm response registration. Each session began with on-screen instructions prompting the participant to rest their response-hand on the homebase to standardize movement onset position across trials.

### 2.4. Upper Extremity Kinematics Data Acquisition & Processing

We recorded upper extremity movements at 240 Hz using an optical motion capture system (Optitrack Prime 41 cameras, Motive v2.1 software, NaturalPoint Inc., Corvallis, OR, USA) and analyzed kinematic data using a custom MATLAB 2024b script (Mathworks Inc., Natick, MA, USA). We placed 30 reflective markers on participants’ upper extremities (11 markers on each upper extremity) and head (8 markers) [38], and used the response-hand lateral wrist marker (“wrist-out”) as the primary marker to quantify reach kinematics. After placing these markers, the Motive::Body software generated arm skeletons and a head rigid body to track movements, as well as rigid bodies for the touchscreen monitor (using four corner markers) and central homebase (using four equally spaced markers) to ensure a constant touchscreen-to-homebase distance. Marker trajectories were synchronized with Presentation task events via LSL. For each trial, we segmented wrist-out trajectories into three time-windows: (1) the cue interval (from arrow cue onset to 40 msec after), (2) the pre-recall array interval (from 40 msec before to recall array onset), and (3) the reach interval (from 100 msec before recall array onset to touchscreen response onset). We calculated 3D velocity using a Savitzky-Golay smoothing filter (17-point window, 5^th^ order polynomial) [39], which reduced high-frequency noise while preserving the waveform’s shape, and derived instantaneous speed as the Euclidean norm of the velocity vector, where *v_x_(t), v_y_(t), v_z_(t)* represent the velocities in the x, y, and z directions at time *t*, respectively:

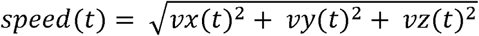

We applied four kinematic quality control criteria prior to further behavioral (see *section 2.5*) and EEG (see *section 2.6*) analyses to ensure reliable reach detection and confirm that movements were initiated in response to the recall array rather than prematurely or due to noise: (1) We verified that the participants’ wrist remained stationary after arrow cue onset by confirming that wrist-out speed during the cue interval had a narrow range of max – min speed < 0.03 m/sec (cue speed stationarity = 1); (2) We applied the same stationarity threshold to the pre-recall array interval to ensure the wrist remained stationary just before recall array onset (recall array speed stationarity = 1); (3) We defined movement onset as the final time point prior to peak speed at which wrist-out speed fell below a within-subject dynamic threshold (i.e., 5 x the SD of baseline cue interval speed). We retained trials only if a valid movement onset time was found before the touchscreen response onset (movement onset = 1); and (4) We verified that movement onset occurred after the recall array onset, ensuring that participants initiated movement in response to the recall array onset rather than prematurely (true pre-reach duration found = 1).

### 2.5. Behavioral Data Processing & Analysis

For each valid memory and motor-only trial (see *section 2.4*), we extracted the following behavioral measures: (1) Error distance (ED; cm), (2) robust coefficient of variation (%), (3) accuracy (%), (4) normalized ED, (5) reaction duration (sec), (6) pre-reach duration (sec), (7) reach duration (sec), (8) reach trajectory length (m), and (9) reach speed (m/sec). We defined ED as the Euclidean distance between the target and index finger response x, y coordinates on the touchscreen monitor:

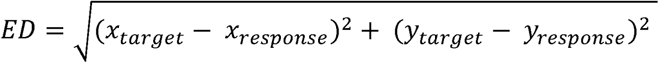

The continuous measure of ED, which we converted from pixel to cm, reflects spatial precision in WM performance, with smaller value indicating better WM performance. We designed the task to produce a continuous rather than binary performance measure to better capture real-world memory performance, such as estimating the approximate location of objects in space rather than simply recalling their presence or absence. To better understand the dispersion of EDs, accounting for the non-normal distribution of the ED data, we then calculated the robust (non-parametric) coefficient of variation (rCV), where rCV is interquartile range divided by median ED. To calculate accuracy, we defined ‘correct’ trials as those with ED shorter than within-subject threshold value, which is calculated as 2 SD above each participant’s mean motor-only ED, and we divided the ‘correct’ trials by all trials for each participant. Therefore, each participant had a unique threshold value. Further, to better isolate the cognitive component of task performance by minimizing motor-related interference, we normalized memory trial performance with motor-only trial performance for each participant:

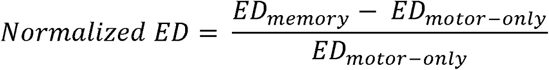

We defined reaction duration to be the latency between the recall array onset and touchscreen response onset. We then divided this into pre-reach duration and reach duration. Pre-reach duration is the latency between the recall array onset and movement onset, while reach duration is the latency between movement onset and touchscreen response onset. We defined the reach trajectory length to be the total 3D wrist-out path length from movement onset to touchscreen response onset, computed by summing the Euclidean distances between successive time points across the x, y, and z dimensions, where (*x_i_, y_i_, z_i_*) is the wrist position at time point *i*, and *N* is the number of sampled time points:

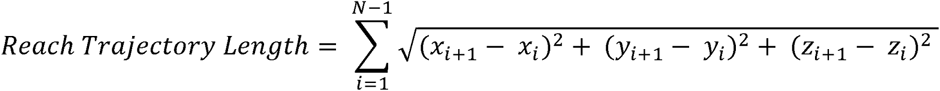

We defined reach speed as trajectory length divided by reach duration. For quality control and artifact detection, we additionally generated 3D wrist trajectory plots and velocity/speed time series for each trial, with manual inspection interface to review kinematics and identify tracking anomalies. Because target location inherently affects reach time and trajectory length, we prioritized reach speed over these individual measures for motor-related analyses.

### 2.6. EEG Activity Processing & Analysis

We recorded continuous EEG using a 128-electrode configuration BioSemi ActiveTwo system (BioSemi Inc., Amsterdam, The Netherlands) at 2,048 Hz. EEG data were streamed via LSL to a computer along with task events and motion capture data streams. We preprocessed the EEG data using custom MATLAB scripts incorporating EEGLAB (v2024.1) and FieldTrip (20250114 release). EEG data were filtered using the zero-phase Chebyshev Type II filter (“filtfilt” function in MATLAB, passband ripple “Apass” = 1 dB, and stopband attenuation “Astop” = 65 dB) [27, 33] and down-sampled from 2,048 Hz to 512 Hz. Next, “bad” electrodes were detected based on kurtosis, probability, and spectral deviation criteria using a ± 3 SD threshold, separately across three metrics. These identified “bad” electrodes were removed and interpolated based on neighboring electrodes, using spherical interpolation. All the electrodes were re-referenced offline to a common average reference. External electrodes were excluded prior to re-referencing, and a temporary reference electrode was inserted and removed to allow proper average referencing (“pop_reref” function in EEGLAB). It has been shown that 1 – 2 Hz high-pass filtered EEG data yield the optimal independent component analysis (ICA) decomposition results in terms of signal-to-noise ratio [40–43]. Therefore, after running the Infomax ICA (“runica” function in EEGLAB, the number of retained principal components matched the rank of the EEG data) on 2-40 Hz bandpass-filtered data and obtaining the decomposition matrices (weight and sphere matrices), these matrices were transferred and applied to 0.01 – 40 Hz bandpass-filtered data. We used a conservative high-pass filter (0.1 Hz) based on the evidence that high-pass filters ≤ 0.1 Hz introduce fewer artifacts into ERP waveforms [42, 44]. We prioritized the use of ICA to identify and remove eye movement artifacts. ICs were labeled using the ICLabel algorithm [45]. ICs with summed probabilities > 50% for the specified artifactual IC classes (“Eye,” “Noise,” “Muscle,” “Heart,” “Line Noise,” and “Channel Noise”) were labeled as artifacts and rejected, and the remaining ICs were back-projected to the sensor space. The resulting EEG data was split into temporal epochs. For all load types (i.e., 1- to 4-disc), memory epochs were time-locked to the encoding array onset (time 0), spanning from -200 to 1,400 msec (i.e., pre- to post-encoding array onset). Epochs were baseline-corrected relative to the pre-encoding array onset interval from -200 to 0 msec to remove low frequency offsets. Epochs were rejected using a trial-wise maximum voltage criterion based on median absolute deviation (MAD) of the maximum absolute voltage per epoch. The rejection threshold was defined adaptively per participant as the smallest value between 3 – 15 MADs that removed fewer than 20% of trials.

By load type, the final thresholds (mean ± SD) were 3.2 ± 0.5 for 1-disc, 3.2 ± 0.9 for 2-disc, 3.5 ± 1.2 for 3-disc, and 3.2 ± 0.5 for 4-disc. By target hemifield across load types, the final thresholds for left hemifield were 3.1 ± 0.3 for 1-disc, 3.3 ± 0.9 for 2-disc, 3.7 ± 1.5 for 3-disc, and 3.4 ± 1.1 for 4-disc and for right hemifield were 3.5 ± 1.0 for 1-disc, 3.3 ± 0.9 for 2-disc, 3.5 ± 1.6 for 3-disc, and 3.3 ± 0.7 for 4-disc. By target hemifield and response-hand, the final thresholds were 3.5 ± 1.2 for LxL (i.e., left target hemifield-left response-hand), 3.4 ± 0.9 for LxR (i.e., left target hemifield-right response-hand), 3.3 ± 0.8 for RxL (i.e., right target hemifield-left response-hand), and 3.3 ± 0.9 for RxR (i.e., right target hemifield-right response-hand). For analyses dividing by both target hemifield and response hand (i.e., condition) across load types, we retained data only when rejection thresholds did not exceed 15 MADs per condition-load combination. This resulted in including only 22 participants for LxL, 21 for LxR, 23 for RxL, and 23 for RxR (see *section 3.3.4*).

Only trials that passed all quality control steps (see *section 2.4*) were included in the analysis. ERPs were computed for each load condition by averaging across trials for each participant. Guided by prior literature on WM-related ERPs at posterior sites using lateralized tasks, we defined 300–900 msec post-encoding array onset as the *a priori* window for averaging ERP amplitudes during the delay period amplitudes [3, 9, 46]. For each participant, we extracted amplitudes from several electrode combinations. In addition to the PO7/8 pair, which has shown to be the most representative pair for examining contralateral delay activity [9, 47], we used the average of three posterior electrode pairs including PO7/8, P7/8, and PO3/4, as these sites have also been frequently employed in prior WM-related work [9, 48]. For load-related contralateral activity, ipsilateral activity, and CDA (i.e., contralateral – ipsilateral difference activity), we applied the same *a priori* 300–900 msec window and the same electrode combinations to extract mean amplitudes from each participant. To explore load-related differences in the early visual-evoked potential, the mean N1 amplitude from each participant was measured between 150–250 msec at posterior sites. Given the lateralized nature of our task, we focused on posterior site(s) contralateral to the target hemifield, where N1 activity is typically strongest [46, 49–51]. The time window and electrode locations were selected based on past literature [52–54] and confirmed and adjusted by inspecting the timing and topography of the major voltage fluctuations in the grand averages. For simplicity, from here on, we refer to the early visual-evoked N1 period (150–250 msec) as *interval 1* and the subsequent delay period (300–900 msec) as *interval 2*. We used the mean amplitudes within these pre-defined time intervals for subsequent statistical analyses.

### 2.7. Statistical Analyses

#### 2.7.1. Behavioral Performance Analyses

First, to examine the differences in the defined behavioral measures by load (motor-only (if applicable), 1-disc, 2-disc, 3-disc, 4-disc), we conducted repeated-measures analyses of variance (ANOVAs) with Load as a within-subject factor for each behavioral measure. Second, to examine the differences in representative behavioral measures by target hemifield (left, right) across memory loads, we conducted two-way repeated-measures ANOVA with Target Hemifield and Load as within-subject factors for each behavioral measure. Third, to examine the differences in representative behavioral measures by response-hand (left, right) across memory loads, we conducted two-way repeated-measures ANOVAs with Response-Hand and Load as within-subject factors for each behavioral measure. Finally, to examine the differences in representative behavioral measures by different conditions (i.e., four different combinations of target hemifield and response-hand) across memory loads, we conducted two-way repeated-measures ANOVAs with Condition (LxL, LxR, RxL, RxR) and Load as within-subject factors. For all analyses, when appropriate, Greenhouse-Geisser correction was used to account for violation of the assumption of sphericity. For post-hoc analysis, we used Bonferroni adjustment, and we reported effect sizes as partial eta squared (η*_p_*^2^). To note, for all analyses involving Response-Hand, we excluded two participants who are left-hand dominant and one participant who had usable data from only one hand after quality check. This ensured a more coherent framework for examining the effects of handedness on behavioral performance, resulting in a final sample of 23 participants for these analyses.

#### 2.7.2. EEG Activity Analyses

First, to examine the differences in the defined parieto-occipital site activity (PO7, Oz, PO8) by load (1-disc to 4-disc), we conducted repeated-measures ANOVAs with Load as a within-subjects factor for the mean activity amplitudes at each electrode at both intervals 1 and 2, separately. Second, to examine the lateralized differences in the *a priori* parieto-occipital site activity by load, we averaged the mean amplitudes of contralateral activity of left and right target hemifield trials, averaged the mean amplitude of ipsilateral activity of left and right target hemifield trials, and calculated the CDA as the difference between the mean contralateral activity and the mean ipsilateral activity. For left target hemifield trials, we measured contralateral activity from right posterior site(s), and for right target hemifield trials, from left posterior site(s). Analyses focused first on the PO7/PO8 pair and then on the average of three posterior electrode pairs (PO7/8, P7/8, PO3/4). We conducted repeated-measures ANOVAs with Load as a within-subjects factor for the mean contralateral activity amplitudes during interval 1 using the PO7/PO8 pair and the average of three posterior electrode pairs, separately. We then conducted repeated-measures ANOVAs with Load as a within-subjects factor for the mean contralateral, ipsilateral, and CDA amplitudes during interval 2 using the PO7/PO8 pair and the average of three posterior electrode pairs, separately. Third, to examine the differences in the defined parieto-occipital site activity (PO7, Oz, PO8) by different conditions (LxL, LxR, RxL, RxR), we conducted repeated-measures ANOVAs with Condition as a within-subjects factor for the mean activity amplitudes at each electrode at both intervals 1 and 2, separately. Finally, to examine the differences in representative ERP component measures (contralateral N1, CDA) by response-hand across loads, we conducted two-way repeated-measures ANOVAs with Response-Hand (left, right) and Load as within-subjects factors for the mean contralateral activity amplitudes for interval 1 and mean CDA amplitudes for interval 2, separately. We averaged the LxL and RxL data for left response-hand, and LxR and RxR data for right response-hand. For all analyses, when appropriate, Greenhouse-Geisser correction was used to account for violation of the assumption of sphericity. For post-hoc analysis, we used Bonferroni adjustment, and we reported effect sizes as partial eta squared (η*^p^*^2^). Again, for analyses involving Response-Hand, we only included 23 right-hand dominant participants who have both left and right response-hand data.

#### 2.7.3. Exploratory Whole Brain Analyses

In addition, because analyses restricted to hypothesis-driven *a priori* electrodes may overlook broader effects, we performed exploratory statistical cluster plots (SCPs) [33, 34, 55] across all electrodes and time points for each load pair to more fully characterize the spatiotemporal distribution of load-related activity. For each participant and electrode, we calculated the mean ERP difference between memory loads by subtracting the within-subject mean waveform at the lower load from the corresponding mean waveform at the higher load.

We then performed paired sample t-tests on these difference waveforms across all participants at each electrode-timepoint pair. To correct for multiple comparisons, we applied cluster-based permutation testing (5,000 permutations, Monte Carlo method), using dependent-samples t-tests as the test statistic, a two-sided significance level of α = 0.05 (tail probabilities corrected accordingly), and the weighted cluster mass statistic [33, 56] with a parametric cluster significance level of α = 0.05. Clusters were defined spatiotemporally based on electrode neighborhood information from the BioSemi 128 layout (triangulation method) and temporal adjacency, with a minimum of zero neighboring electrodes required for inclusion. We performed this procedure separately for each load pair, collapsing across all trial types. The resulting intensity plots displayed point-wise t-test results from all 128 electrodes and timepoints for each load comparison (e.g., 2-disc minus 1-disc), providing a compact visualization of the onset and topographical distribution of load-related ERP changes. The x, y, and z axes represent time (msec), electrode location, and the t-statistic value (indicated by color value) at each electrode-timepoint pair. T-statistic values at electrode-timepoint pairs not belonging to any significant cluster were masked (set to zero) and are shown as a teal background.

Statistical analyses (including repeated-measures ANOVAs, topographical scalp maps, and non-parametric SCPs) and plotting of behavioral and EEG activity data were conducted using IBM SPSS Statistics (Version 29.0; IBM Corp., Armonk, NY, USA) and custom MATLAB scripts incorporating EEGLAB (v2024.1) and FieldTrip (20250114 release).

#### 2.7.4. Multivariate Correlation Analyses

To test potential associations between the demographic (age and education) and related standard neuropsychological measurements (MoCA and Digit Span) with the defined task-related behavioral measures, we ran Spearman rank correlations between these variables. Due to the demographic and neuropsychological measures not being normally distributed, all correlations performed in the context of this study used the non-parametric Spearman rank correlation. We similarly ran Spearman rank correlations between representative behavioral measures and mean ERP amplitudes, restricted to matching load conditions. For example, we correlated ED at 1-disc with mean PO7 activity at 1-disc only, not with other unmatched loads.

## 3. RESULTS

### 3.1. Standard Neuropsychological Performances

All 26 participants completed the standard neuropsychological tests of MoCA, which is an assessment to represent global cognitive functioning, and WAIS-IV Digit Span, which is an assessment of WM. Participants scored 27.8 ± 2.1 (mean ± SD; max of 30) on MoCA and 11.5 ± 2.9 (mean ± SD scaled total; max of 19) on WAIS-IV Digit Span.

### 3.2. Behavioral Performances

#### 3.2.1. Behavioral Performance by Load

**Table 1** summarizes the behavioral measures by memory load, and **Figure 2** illustrates both individual-and group-level data for selected measures. We conducted repeated measures ANOVAs with Load (motor-only, 1-disc, 2-disc, 3-disc, 4-disc) as a within-subjects factor for each behavioral measure and found robust load-related effects across multiple measures. Mean ED differed significantly across Loads (*F*(2.40, 59.90) = 185.01, *p* < 0.001, η*^p^*^2^ = 0.88). Post hoc t-tests revealed significant differences between all Load pairs (*p* < 0.05), with ED increasing monotonically as load increased (i.e., from motor-only, 1-disc, 2-disc, 3-disc, to 4-disc) (**Figure 2A**). **Figure 2B** shows individual response distributions relative to the target location (collapsed at the center 0,0) across all disc orientations, with concentric circles marking 1 – 4 cm increments. rCV also differed significantly across Loads (*F*(2.08, 51.95) = 33.76, *p* < 0.001, η*_p_*^2^ = 0.58). Post hoc t-tests revealed significant pairwise differences (*p* < 0.05), with rCV increasing (i.e., greater response dispersion) as load increased, except for motor-only vs. 1-disc (*p* = 1.00), 1-disc vs. 2-disc (*p* = 0.053), and 3-disc vs. 4-disc (*p* = 0.07) (**Figure 2C**). Accuracy, defined using participant-specific thresholds as previously described (group mean threshold = 36.47 pixels; see *section 2.5*), differed significantly across Loads (*F*(1.98, 49.50) = 149.02, *p* < 0.001, η*_p_*^2^ = 0.86). Post hoc t-tests showed significant differences between all Load pairs (*p* < 0.001), with accuracy decreasing as the load increased, except for motor-only vs. 1-disc (*p* = 0.12). Normalized ED, which accounts for motor-related contributions to ED, differed significantly across Loads (*F*(1.75, 43.64) = 119.46, *p* < 0.001, η*_p_*^2^ = 0.83). Post hoc t-tests revealed significant differences between all Load pairs (*p* < 0.001), with normalized ED increasing monotonically as load increased from 1-disc to 4-disc (**Figure 2D**). Reaction duration differed significantly across Loads (*F*(1.73, 43.32) = 26.49, *p* < 0.001, η*_p_*^2^ = 0.51). Post hoc t-tests revealed significant differences between all Load pairs (*p* < 0.05), with reaction duration increasing monotonically as load increased from 1-disc to 4-disc, except for 1-disc vs. 2-disc (*p* = 0.066). Pre-reach duration also differed significantly across Loads (*F*(1.62, 40.39) = 48.21, *p* < 0.001, η*_p_*^2^ = 0.66). Post hoc t-tests revealed significant differences between all Load pairs (*p* < 0.01), with pre-reach duration increasing as load increased from 1-disc to 4-disc (**Figure 2E**). Neither reach duration nor reach trajectory length differed significantly across Loads (all *p* > 0.05). Reach speed differed significantly across Loads (*F*(2.69, 67.13) = 3.70, *p* = 0.019, η*_p_*^2^ = 0.13), but post hoc t-tests did not reveal significant pairwise differences (all *p* > 0.05) (**Figure 2F**).

**Figure 2.**
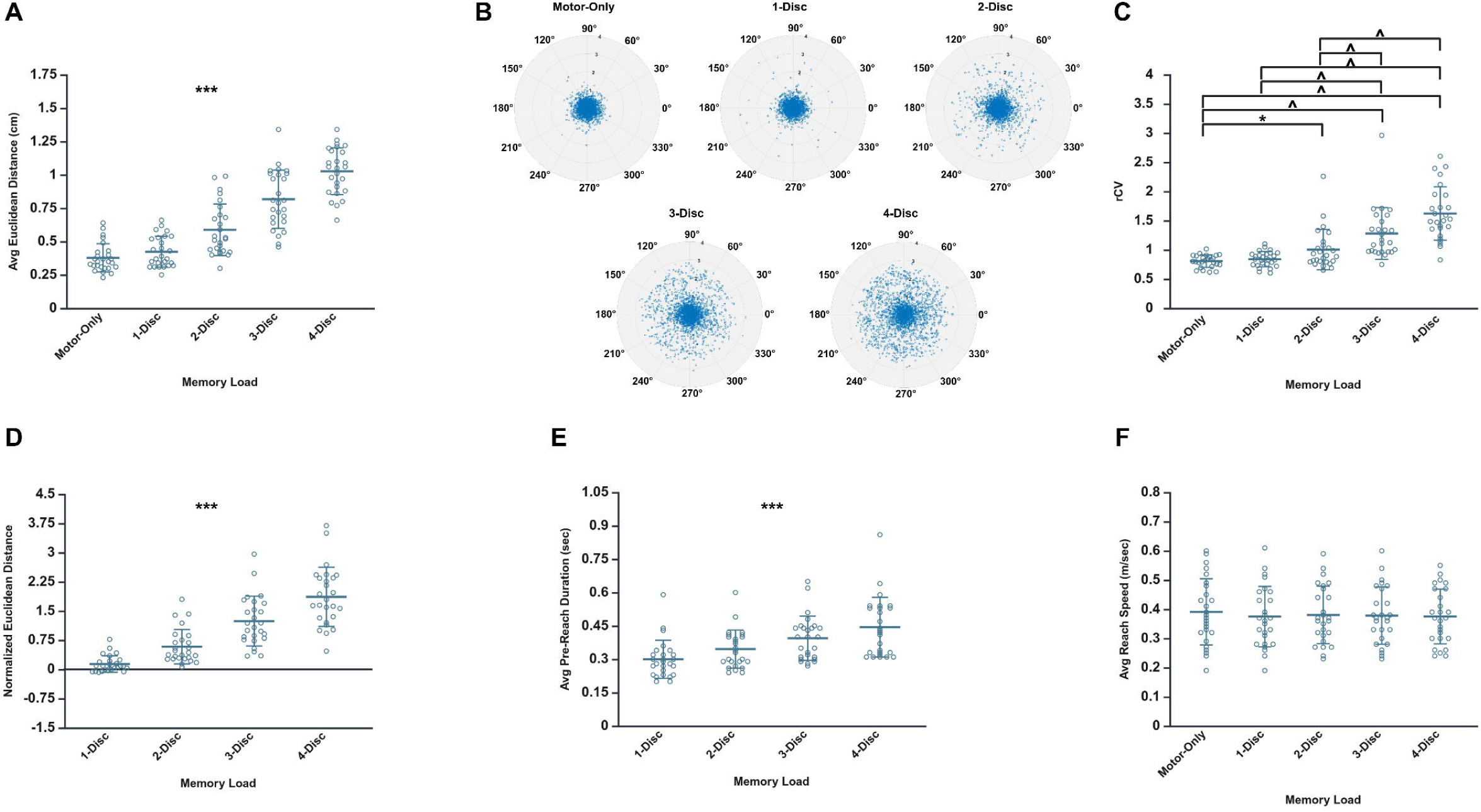
Behavioral Performance Data Across Loads. Unfilled circles show individual participant means for (A) Euclidean distance, (C) robust (non-parametric) coefficient of variation, (D) normalized Euclidean distance, (E) pre-reach duration, and (F) reach speed. The central line and error bars indicate group means +/-SD. (B) shows individual participant’s response locations from all 26 participants plotted relative to the target location, which is collapsed at the center (0,0). Data are shown across all disc opening orientations (in degrees) and all included trials, separated by memory load to illustrate the dispersion of responses. Concentric circles represent radial distances from the target in 1cm increments, ranging from 1cm to 4cm. Values beyond 4cm are not shown. * indicates *p* < 0.05. ^ indicates *p* < 0.001. *** indicates that all load interactions are statistically significant (*p* < 0.05).

#### 3.2.2. Behavioral Performance by Load and Target Hemifield

**Supplementary Table 1** summarizes representative behavioral measures by memory load and target hemifield. We conducted two-way repeated-measures ANOVAs with Target Hemifield (left, right) and Load (motor-only, 1-disc, 2-disc, 3-disc, 4-disc) as within-subjects factors for each behavioral measure to compare performance across target hemifields and load types. Mean ED differed significantly across Loads (*F*(2.34, 58.54) = 187.22, *p* < 0.001, η*_p_*^2^ = 0.88). Post hoc t-tests replicated the pattern reported in *section 3.2.1*, with all Load pairs differing significantly (*p* < 0.05) and ED increasing with load. Neither the main effect of Target Hemifield nor the Load x Target Hemifield interaction was significant (all *p* > 0.05). rCV differed significantly between Target Hemifields (*F*(1.00, 25.00) = 4.95, *p* = 0.035, η*_p_*^2^ = 0.17), with greater response dispersion for targets in the right-hemifield than left-hemifield. rCV also differed significantly across Loads (*F*(1.88, 47.08) = 26.02, *p* < 0.001, η*_p_*^2^ = 0.51). Post hoc t-tests again replicated the pattern reported in *section 3.2.1*, with rCV increasing as load increased (*p* < 0.05), except between motor-only vs. 1-disc, 1-disc vs. 2-disc, and 3-disc vs. 4-disc. The Load x Target Hemifield interaction was not significant (*F*(2.45, 61.27) = 2.92, *p* = 0.051, η*_p_*^2^ = 0.11).

Pre-reach duration differed significantly across Loads (F(1.59, 39.83) = 48.05, *p* < 0.001, η*_p_*^2^ = 0.66). Post hoc t-tests replicated the pattern reported in *section 3.2.1*, with all Load pairs differing significantly (*p* < 0.01) and pre-reach duration increasing as load increased. Neither the main effect of Target Hemifield nor the Load x Target Hemifield interaction was significant (all *p* > 0.05). Reach speed differed significantly across Loads (*F*(2.56, 63.93) = 3.96, *p* = 0.016, η*_p_*^2^ = 0.14), but as in *section 3.2.1*, post hoc t-tests did not reveal significant pairwise differences (all *p* > 0.05). Neither the main effect of Target Hemifield nor the Load x Target Hemifield interaction was significant (all *p* > 0.05).

#### 3.2.3. Behavioral Performance by Load and Response-Hand

**Supplementary Table 2** summarizes representative behavioral measures by memory load and response-hand. As noted in *section 2.7*, this analysis included 23 right-hand dominant participants who have both left and right response-hand data to provide a coherent framework for examining handedness effects on behavioral performance. We conducted two-way repeated-measures ANOVAs with Response-Hand (left, right) and Load (motor-only, 1-disc, 2-disc, 3-disc, 4-disc) as within-subjects factors for each behavioral measure to compare performance across response-hands and load types. Mean ED differed significantly across Loads (*F*(2.54, 55.88) = 195.76, *p* < 0.001, η*_p_*^2^ = 0.90). Post hoc t-tests replicated the pattern reported in *section 3.2.1*, with all Load pairs differing significantly (*p* < 0.001), except for motor-only vs. 1-disc (*p* = 0.061). Neither the main effect of Response-Hand nor the Load x Response-Hand interaction was significant (all *p* > 0.05). rCV differed significantly across Loads (*F*(1.82, 39.94) = 32.41, *p* < 0.001, η*_p_*^2^ = 0.60). Post hoc t-tests revealed that rCV significantly increased with load (*p* < 0.01), except between motor-only vs. 1-disc, motor-only vs. 2-disc, 1-disc vs. 2-disc, and 3-disc vs. 4-disc (all *p* > 0.05). Neither the main effect of Response-Hand nor the Load x Response-Hand interaction was significant (all *p* > 0.05). Pre-reach duration differed significantly between Response-Hands (*F*(1.00, 22.00) = 4.32, *p* = 0.049, η*_p_*^2^ = 0.16), with longer pre-reach durations for the right (dominant) hand than the left hand. Pre-reach duration also differed significantly across Loads (*F*(1.60, 35.29) = 40.65, *p* < 0.001, η*_p_*^2^ = 0.65). Post hoc t-tests replicated the pattern reported in *section 3.2.1*, with all Load pairs differing significantly (*p* < 0.01) and pre-reach duration increasing with load. The Load x Response-Hand interaction was not significant (*F(*2.06, 45.39) = 1.68, *p* = 0.198, η*_p_*^2^ = 0.07). Reach speed differed significantly between Response-Hands (*F*(1.00, 22.00) = 4.55, *p* = 0.044, η*_p_*^2^ = 0.17), with faster reaches using the right (dominant) hand than the left hand. Neither the main effect of Load nor the Load x Response-Hand interaction was significant (all *p* > 0.05).

#### 3.2.4. Behavioral Performance by Load and Condition (Target Hemifield and Response-Hand Combination)

**Supplementary Table 3** summarizes representative behavioral measures by memory load and condition, which includes four combinations of Target Hemifield and Response-Hand (LxL, LxR, RxL, RxR). As noted in *section 2.7*, this analysis includes 23 right-hand dominant participants who have both left and right response-hand data. We conducted two-way repeated-measures ANOVAs with Condition (LxL, LxR, RxL, RxR) and Load (motor-only, 1-disc, 2-disc, 3-disc, 4-disc) as within-subjects factors for each behavioral measure to compare performance across conditions and load types. Mean ED differed significantly across Loads (*F*(2.50, 55.02) = 201.63, *p* < 0.001, η*_p_*^2^ = 0.90). Post hoc t-tests replicated the pattern reported in *section 3.2.3*. Neither the main effect of Condition nor the Load x Condition interaction was significant (all *p* > 0.05). rCV differed significantly across Conditions (*F*(3.00, 66.00) = 2.83, *p* = 0.045, η*_p_*^2^ = 0.11). Post hoc t-tests revealed greater dispersion in RxL compared to LxR (*p* = 0.036), indicating that variability increased when participants used the non-dominant hand for contralateral targets relative to when they used the dominant hand for contralateral targets. rCV also differed significantly across Loads (*F*(1.78, 39.26) = 34.57, *p* < 0.001, η*_p_*^2^ = 0.61). Post hoc t-tests replicated the pattern reported in *section 3.2.3*. The Load x Condition interaction was not significant (*F*(4.77, 104.92) = 0.976, *p* = 0.433, η*_p_*^2^ = 0.04). Pre-reach duration differed significantly across Loads (*F*(1.61, 35.51) = 41.24, *p* < 0.001, η*_p_*^2^ = 0.65). Post hoc t-tests replicated the pattern reported in *section 3.2.3*. Neither the main effect of Condition nor the Load x Condition interaction was significant (all *p* > 0.05). Reach speed differed significantly across Conditions (*F*(1.37, 30.21) = 4.70, *p* = 0.028, η*_p_*^2^ = 0.18). Post hoc t-tests revealed faster reach speeds in RxR compared to RxL (*p* = 0.021), indicating faster reaches using the dominant hand than the non-dominant hand for targets in the right hemifield. Neither the main effect of Load nor the Load x Condition interaction was significant (all *p* > 0.05).

### 3.3. EEG Activity

#### 3.3.1. ERP Changes as a Function of Load

We examined load-related EEG activity (1-disc to 4-disc) pooled across all participants, time-locked to the encoding array onset, and collapsed across target hemifields and response-hand. To demonstrate the overall distribution of the electrophysiological response, **Figure 3** shows representative ERPs from key scalp sites. The waveforms displayed typical morphology, with an early visual-evoked N1 (150 – 250 msec; interval 1) followed by sustained delay-period (300 – 900 msec; interval 2) activity. At parieto-occipital sites (PO7, Oz, PO8), higher loads evoked greater negativity during interval 1 and extended into interval 2. Frontal (Fz) and central (Cz) sites showed the opposite pattern, with relative positivity at higher loads. Repeated-measures ANOVAs for the mean activity amplitude at each parieto-occipital site during interval 1 confirmed significant main effects of Load for all three sites (**Table 2**; all *p* < 0.01), with negativity increasing as load increased. Post hoc t-tests revealed significant differences in PO7 for 1-disc vs. 3-disc, 1-disc vs. 4-disc, and 3-disc vs. 4-disc; in Oz for 1-disc vs. 4-disc, 2-disc vs. 4-disc, and 3-disc vs. 4-disc; and in PO8 for 1-disc vs. 2-disc, 1-disc vs. 3-disc, and 1-disc vs. 4-disc (all *p* < 0.05). Repeated-measures ANOVAs for each parieto-occipital site during interval 2 also confirmed significant main effects of Load for Oz and PO8 (**Table 2**; all *p* < 0.01), with negativity increasing as load increased. Post hoc t-tests revealed significant differences in Oz for 1-disc vs. 4-disc, and in PO8 for 1-disc vs. 3-disc and 1-disc vs. 4-disc (all *p* < 0.05). PO7 did not show significant Load effects during interval 2.

**Figure 3.**
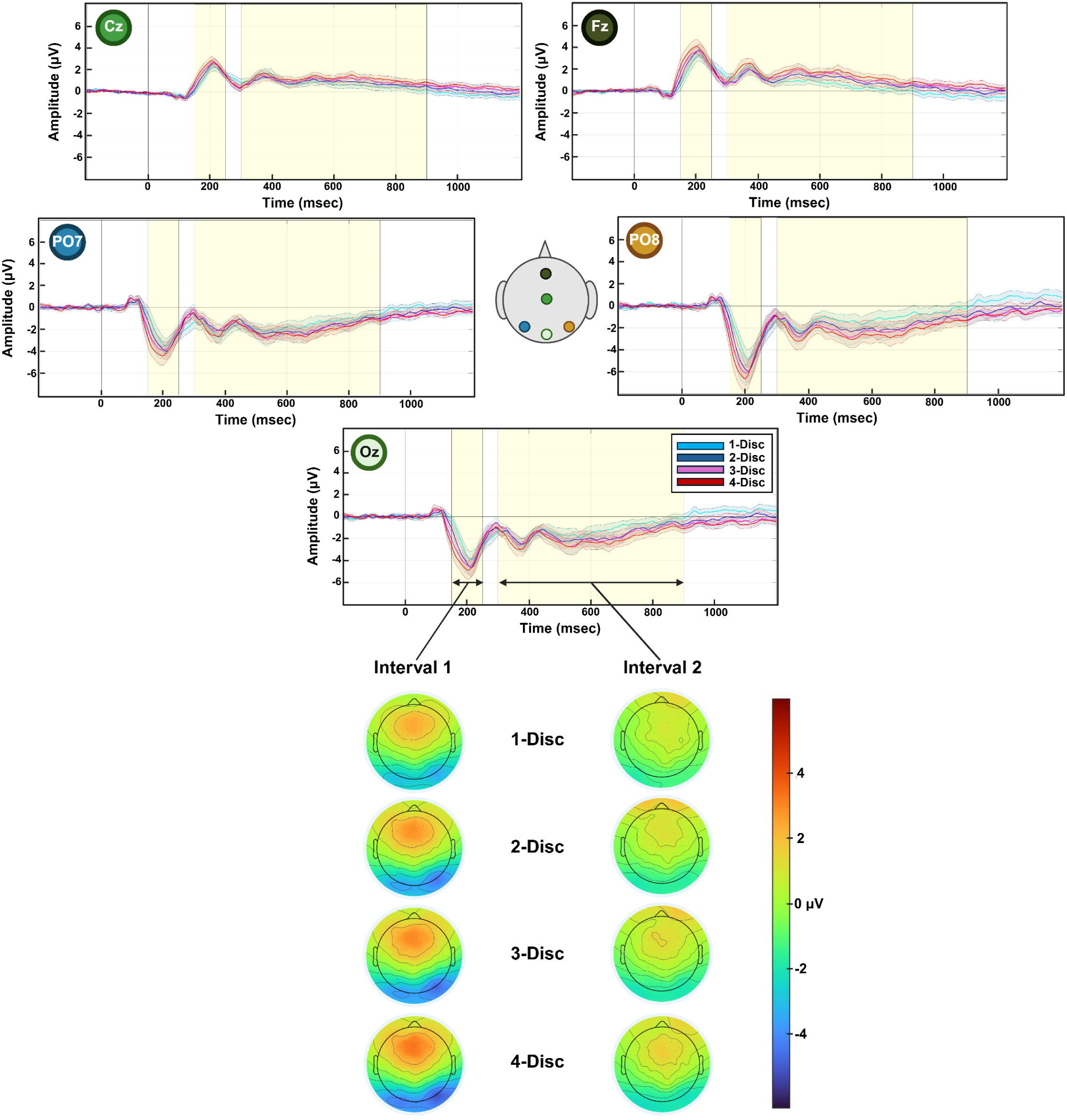
Neurophysiological Activities Across Loads. The top panel shows average waveforms with SD in colored shades time-locked to the encoding array onset (time 0) across different memory loads. Highlighted regions correspond to the intervals selected for preplanned statistical analysis. Colored circles indicate scalp location for each corresponding waveform. The bottom panel shows topographical scalp maps displaying mean amplitude at the highlighted intervals (interval 1 = 150 – 250 msec; interval 2 = 300 – 900 msec).

Because analyses at electrode locations selected *a priori* can miss broader effects, we performed exploratory SCPs across all electrodes and time points for each load pair (see *section 2.7.2*) to examine load-related effects (**Figure 4**). These intensity plots show the results of paired t-tests comparing different load pairs (e.g., 2-disc minus 1-disc). Compared to the baseline memory load (1-disc), 2-disc minus 1-disc resulted in only posterior negativity during interval 2. 3-disc minus 1-disc showed widespread frontal and central positivity during both intervals and posterior negativity during interval 2 only. 4-disc minus 1-disc showed an even stronger effect with widespread frontal and central positivity and posterior negativity during both intervals. 4-disc minus 2-disc resembled the 4-disc minus 1-disc pattern but with weaker effects, particularly during interval 2, with the loss of posterior negativity and weaker frontal positivity. 4-disc minus 3-disc and 3-disc minus 2-disc generated no significant clusters during both intervals.

**Figure 4.**
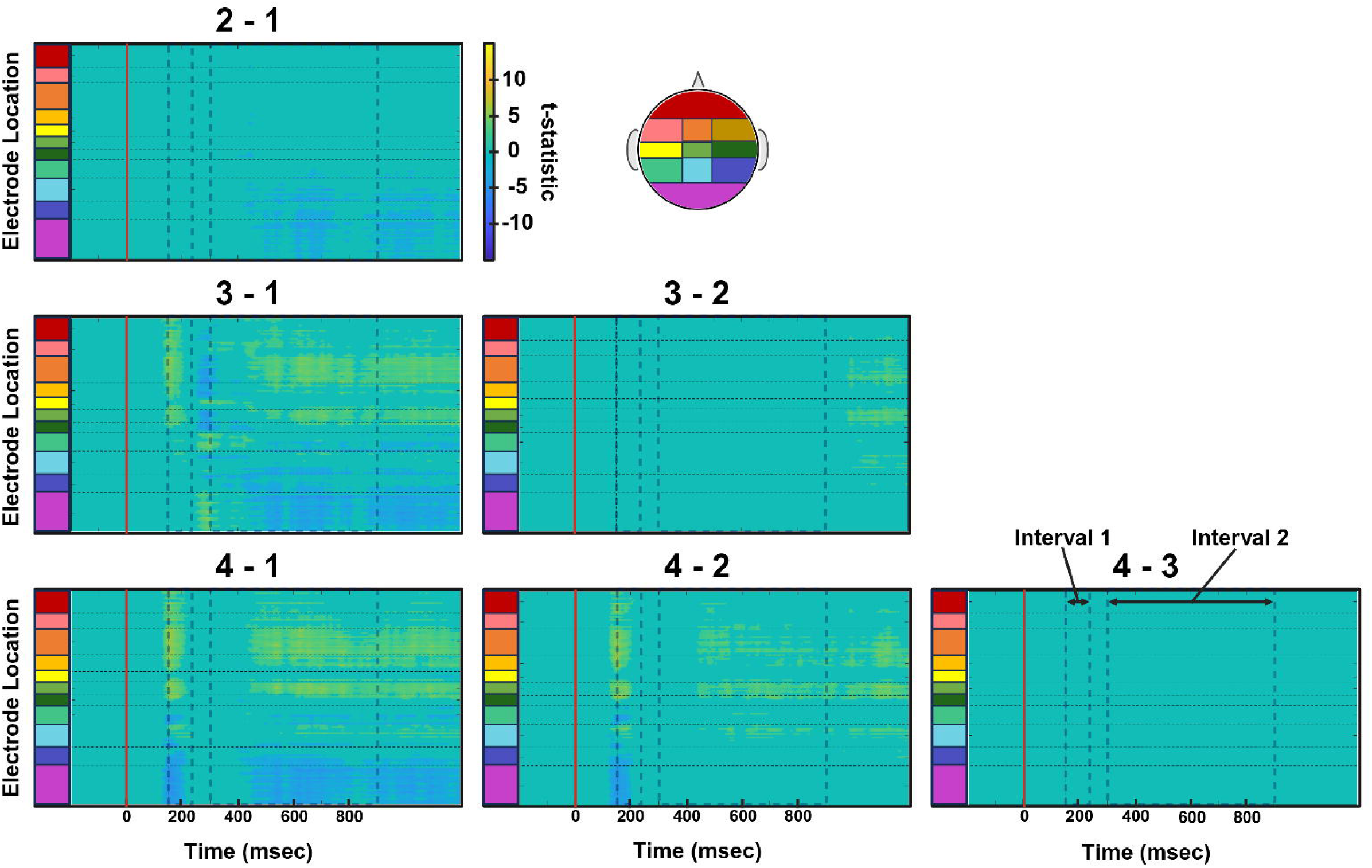
Load-Related Statistical Cluster Plots. Data used in analyses are time-locked to the encoding array onset (red vertical line) with a 200 msec pre-encoding array onset baseline. Color values indicate the t-statistic result of pointwise t-tests comparing different load pairs as indicated by the numbers above each plot (e.g., 2 - 1 = 2-disc minus 1-disc) conducted at each time point (x-axis) and electrode locations (y-axis). Spatiotemporal clusters illustrate regions and intervals of relative positivity (yellow) or negativity (blue) in task-evoked potentials recorded during simultaneous higher load (with respect to lower load pair).

#### 3.3.2. ERP Changes as a Function of Load and Target Hemifield

We next examined load-related EEG activity by target hemifield to assess contralateral activity, ipsilateral activity, and CDA. For left target hemifield trials, we measured contralateral activity from right posterior site(s), and for right target hemifield trials, from left posterior site(s). Analyses focused first on the PO7/PO8 pair (**Figure 5**) and then on the average of three posterior electrode pairs (PO7/8, P7/8, PO3/4; Figure 6).

**Figure 5.**
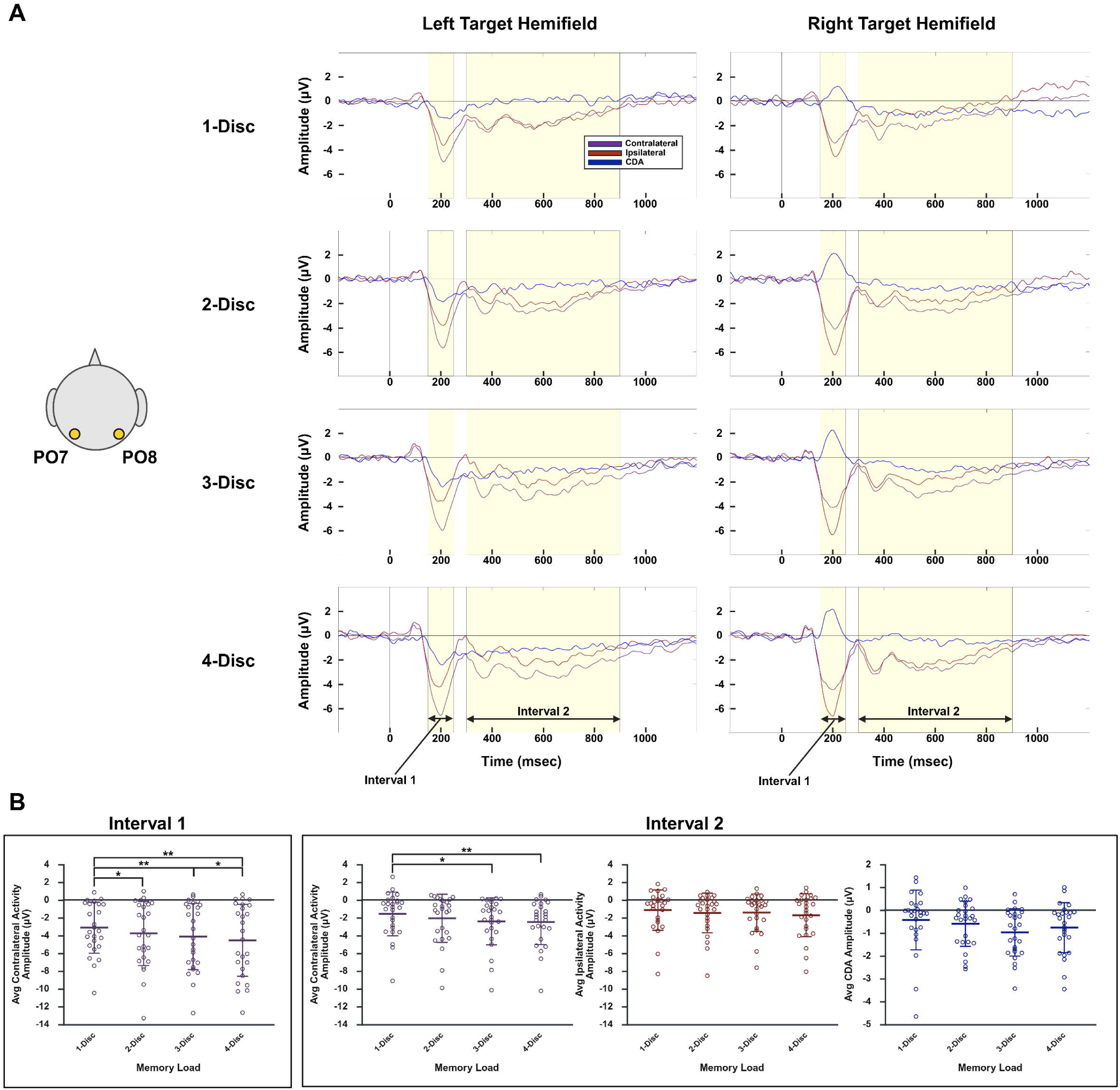
Neurophysiological Activities (Contralateral, Ipsilateral, CDA) Across Loads Using PO7/PO8 Pair. (A) shows average contralateral (purple), ipsilateral (red), and CDA (i.e., contralateral minus ipsilateral difference; blue) waveforms time-locked to the encoding array onset (time 0) across different memory loads. Highlighted regions correspond to the intervals selected for preplanned statistical analysis. For left target hemifield, we probed contralateral activity from PO8, and for right target hemifield, we probed contralateral activity from PO7. In (B), unfilled circles show individual participant means for contralateral activity (purple) during interval 1 (left panel) and for contralateral activity (purple), ipsilateral activity (red), and CDA (blue) during interval 2 (right panel). The central line and error bars indicate group means +/- SD. * indicates *p* < 0.05. ** indicates *p* < 0.01.

**Figure 6.**
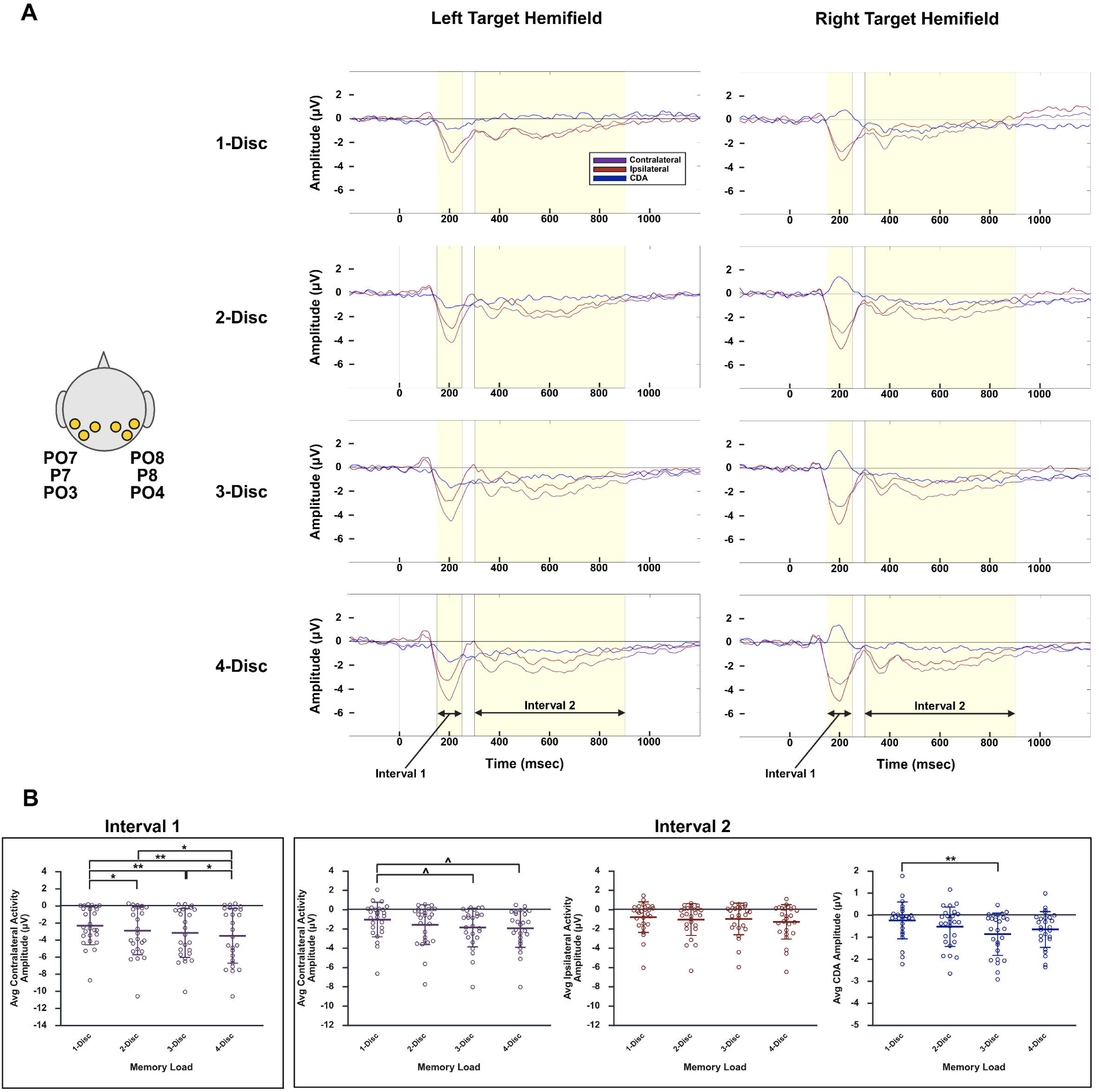
Neurophysiological Activities (Contralateral, Ipsilateral, CDA) Across Loads Using 3 Posterior Electrode Pairs. (A) shows average contralateral (purple), ipsilateral (red), and CDA (i.e., contralateral minus ipsilateral difference; blue) waveforms time-locked to the encoding array onset (time 0) across different memory loads. Highlighted regions correspond to the intervals selected for preplanned statistical analysis. For left target hemifield, we probed contralateral activity from the average of PO8, P8, and PO4, and for right target hemifield, we probed contralateral activity from the average of PO7, P7, and PO3. In (B), unfilled circles show individual participant means for contralateral activity (purple) during interval 1 (left panel) and for contralateral activity (purple), ipsilateral activity (red), and CDA (blue) during interval 2 (right panel). The central line and error bars indicate group means +/- SD. * indicates *p* < 0.05. ** indicates *p* < 0.01. ^ indicates *p* < 0.001.

**Figure 5A** shows contralateral, ipsilateral, and CDA waveforms for each load by target hemifield using the PO7/PO8 pair. Averaging across hemifields, repeated-measures ANOVAs for the mean contralateral activity amplitude confirmed significant main effects of Load during both intervals 1 and 2 (**Table 2**; all *p* < 0.001), with negativity increasing as load increased. During interval 1, post hoc t-tests revealed significant differences between 1-disc vs. 2-disc, 1-disc vs. 3-disc, 1-disc vs. 4-disc, and 3-disc vs. 4-disc (all *p* < 0.05) (**Figure 5B, left panel**). During interval 2, post hoc t-tests revealed significant differences between 1-disc vs. 3-disc and 1-disc vs. 4-disc (all *p* < 0.05) (**Figure 5B, right panel**). Repeated-measures ANOVA for the mean ipsilateral activity amplitude during interval 2 also showed significant main effects of Load, but post hoc t-tests did not reveal significant pairwise differences (all *p* > 0.05) (**Table 2**; **Figure 5B, right panel**). Despite showing increasing negativity with load up to 3-disc, the mean CDA amplitude during interval 2 did not differ significantly across Loads (**Table 2**; **Figure 5B, right panel**).

Similarly, **Figure 6A** shows contralateral, ipsilateral, and CDA waveforms for each load by target hemifield using the average of three posterior electrode pairs. Again, averaging across hemifields, repeated-measures ANOVAs for the mean contralateral activity amplitude confirmed significant main effects of Load during both intervals 1 and 2 (**Table 2**; all *p* < 0.001), with negativity increasing as load increased. During interval 1, post hoc t-tests revealed significant differences between 1-disc vs. 2-disc, 1-disc vs. 3-disc, 1-disc vs. 4-disc, 2-disc vs. 4-disc, and 3-disc vs. 4-disc (all *p* < 0.05) (**Figure 6B, left panel**). During interval 2, post hoc t-tests revealed significant differences between 1-disc vs. 3-disc and 1-disc vs. 4-disc (all *p* < 0.001) (**Figure 6B, right panel**). Repeated-measures ANOVA for the mean ipsilateral activity amplitude during interval 2 also showed significant main effects of Load, but post hoc t-tests did not reveal significant pairwise differences (all *p* > 0.05) (**Table 2**; **Figure 6B, right panel**). Repeated-measures ANOVA for the mean CDA amplitude during interval 2 confirmed significant main effects of Load, with increasing negativity as load increased up to 3-disc (**Table 2**). Post hoc t-tests revealed significant difference between 1-disc vs. 3-disc (*p* = 0.003) (**Figure 6B, right panel**).

#### 3.3.3. ERP Changes as a Function of Condition (Target Hemifield and Response-Hand Combination)

Because using the non-dominant hand may increase task demands, we examined EEG activity by four combinations of Target Hemifield and Response-Hand (i.e., LxL, LxR, RxL, RxR). This analysis included 23 right-hand dominant participants who have both left and right response-hand data, time-locked to encoding array onset and collapsed across memory loads. **Supplementary Figure 1** shows representative ERPs from key scalp sites. These waveforms show clear differences by target hemifield, particularly at lateral posterior sites (PO7, PO8) during interval 2, but little evidence of response-hand effects. Repeated-measures ANOVAs for the mean activity amplitudes at each parieto-occipital site during interval 1 confirmed significant main effects of Condition (LxL, LxR, RxL, RxR) for PO7 and Oz (**Table 3**; all *p* < 0.05), but post hoc t-tests did not reveal significant pairwise differences (all *p* > 0.05). PO8 did not show significant Condition effects during interval 1. Repeated-measures ANOVAs for the mean activity amplitudes at each parieto-occipital site during interval 2 confirmed significant main effects of Condition at PO7 and PO8 (**Table 3**; all *p* < 0.01). Post hoc t-tests did not reveal significant pairwise differences for PO7 (all *p* > 0.05), but for PO8, post hoc t-tests revealed significant differences between LxL vs. RxL (*p* = 0.006), LxR vs. RxL (*p* = 0.001), and LxR vs. RxR (*p* = 0.048), reflecting target hemifield-related effects and stronger differences when the non-dominant hand is used (i.e., LxL vs. RxL) or when target hemifield and response-hand are opposite (i.e., LxR vs. RxL). Oz did not show significant Condition effects during interval 2.

#### 3.3.4. ERP Changes as a Function of Load and Condition

We next examined load-related EEG activity by response-hand to assess contralateral activity, ipsilateral activity, and CDA. For this analysis, we retained the data only if trial rejection thresholds (see *section 2.6*) did not exceed 15 MADs per condition-load combination. For left response-hand, we averaged the LxL and RxL data, where in this case, we measured contralateral activity from right posterior site(s) for LxL and from left posterior site(s) for RxL. Similarly, for right response-hand, we averaged the LxR and RxR data. Analyses focused first on the PO7/PO8 pair and then on the average of three posterior electrode pairs (PO7/8, P7/8, PO3/4).

Using the PO7/PO8 pair (**Supplementary Figure 2; Table 4**), the mean contralateral activity amplitude during interval 1 showed a significant main effect of Load (*F*(2.23, 48.95) = 9.37, *p* < 0.001, η*_p_*^2^ = 0.30), with increasing negativity as load increased. Post hoc t-tests revealed significant differences between 1-disc vs. 3-disc (*p* = 0.011) and 1-disc vs. 4-disc (*p* = 0.001) (**Supplementary Figure 2B, left panel**). Neither the main effect of Response-Hand nor the Load x Response-Hand interaction was significant (all *p* > 0.05). The mean CDA amplitude during interval 2 showed no significant effects of Load, Response-Hand, or their interaction (**Table 4**).

Using the average of three posterior electrode pairs resulted in similar results (**Table 4**). The mean contralateral activity amplitude during interval 1 again showed a significant main effect of Load (*F*(1.95, 42.90) = 11.92, *p* < 0.001, η*_p_*^2^ = 0.35), and post hoc t-tests revealed significant differences between 1-disc vs. 2-disc, 1-disc vs. 3-disc, 1-disc vs. 4-disc, and 3-disc vs. 4-disc (all *p* < 0.05). Neither the main effect of Response-Hand nor the Load x Response-Hand interaction was significant (all *p* > 0.05). The mean CDA amplitude during interval 2 showed no significant effects of Load, Response-Hand, or their interaction (**Table 4**).

### 3.4. Multivariate Correlations

#### 3.4.1. Correlations Between Behavioral Measures and Demographics & Standard Neuropsychological Measures

We examined associations of demographics and standard neuropsychological measures with task-related behavioral measures using Spearman rank correlation analyses (**Table 5**). Since age, education, MoCA, and Digit Span are not normally distributed (Shapiro-Wilk test, *p* < 0.05), we used non-parametric tests. As expected, Digit Span, a standard measure of WM, correlated negatively with ED at 2-disc (*r* = -0.42, *p* = 0.03) and 3-disc (*r* = -0.45, *p* = 0.02), suggesting that higher standard WM performance is associated with better task-related WM performance (i.e., shorter ED) at moderate loads. Digit Span also correlated negatively with rCV at 1-disc (*r* = -0.40, *p* = 0.046), suggesting that higher standard WM performance is associated with less variable task-related WM performance at baseline load. MoCA, a global cognitive measure, also correlated negatively with rCV at 1-disc (*r* = -0.56, *p* < 0.001), suggesting that higher global cognition is associated with less variable task-related WM performance at baseline load. Interestingly, both age and education showed significant negative correlations with reach trajectory length across all loads (**Table 5**), suggesting that older age or higher education level is associated with shorter trajectory paths from homebase to touchscreen response, regardless of loads.

#### 3.4.2. Correlations Between Load-Matched Behavioral and EEG Activity Measures

To explore brain-behavior relationships, we conducted Spearman correlations between behavioral measures and load-matched ERP amplitudes from *section 3.3.2* (**Table 6**). Using the PO7/PO8 pair, ED correlated positively with CDA amplitude during interval 2 at 2-disc (*r* = 0.41, *p* = 0.036), suggesting that better task-related WM performance (i.e., shorter ED) is associated with more negative CDA amplitude at 2-disc. CDA amplitude during interval 2 also correlated positively with reach speed at 2-disc (*r* = 0.51, *p* = 0.008), suggesting that better motor performance (i.e., faster reach speed) is associated with less negative CDA amplitude at 2-disc. Using the average of three electrode pairs, we replicated the positive correlation between reach speed and CDA amplitude at 2-disc (*r* = 0.44, *p* = 0.026) and found no additional significant correlations.

## 4. DISCUSSION

We employed a MoBI system that integrated time-synchronized high-density EEG with 3D motion capture during a touchscreen WM paradigm involving dynamic upper extremity reach and point responses. This approach allowed us to examine how increasing visual WM load shapes behavior and scalp-recorded neural activity when cognitive and motor processes are coupled within a single task, and whether these effects are further modulated by response-hand. By incorporating time-synchronized motion tracking into a WM task, we were able to probe the coordination between cognition and movement through WM-guided actions that better approximate natural behavior, providing a more comprehensive characterization of behavioral performance. Our study addressed that first, increasing load reliably degraded task-related WM performance and slowed motor preparation, as evidenced by larger ED, higher rCV, lower accuracy, and longer pre-reach duration, while leaving core movement execution largely unchanged. Second, parieto-occipital activity tracked load during both the early visual-evoked N1 processing period (150–250 msec) and during the delay period (300–900 msec) but saturated at 3-disc during the delay period only. Posterior-frontal engagement strengthened as load increased. Third, brain-behavior correlation was most evident at a moderate load of 2-discs, where CDA related to both cognitive and motor measures. Finally, overall, handedness exerted selective behavior effects but did not modulate the associated parieto-occipital activities.

### 4.1. Increasing Load Degrades WM Performance and Prolongs Preparation Without Altering Motor Performance

Prior studies have consistently shown that increasing WM load worsens performance, reflecting both task complexity and intrinsic capacity limits [4, 5, 13]. Here, incorporating 3D motion capture during task performance allowed for a more detailed characterization of behavioral outputs, offering greater insight into the cognitive and motor processes engaged by the task. As predicted, increasing memory load degraded task-related WM performance – both ED and normalized ED (i.e., controlled for motor-related contributions to ED) increased monotonically while accuracy declined. The finding that the accuracy of 1-disc was comparable to that of motor-only confirms that 1-disc serves as an effective baseline memory load, with near-ceiling performance across participants. rCV also increased with load, reflecting broader dispersion of responses.

Although most adjacent load comparisons were not significant, the difference between 2-disc and 3-disc was robust, suggesting that higher loads introduce greater uncertainty in WM recall, and that capacity limits likely emerge already at 2-disc; that is, the load at which task-related WM performance in neurotypical adults begins to deteriorate. While rCV did not correlate significantly with EEG activity measures, its sensitivity to load suggests that it may serve as a task-specific behavioral marker of WM capacity. Lower rCV (i.e., less variable WM performance) at 1-disc also showed significant correlation with higher performance in standard WM measures (Digit Span) and global cognitive measures (MoCA), suggesting that more stable performance in baseline memory load reflects higher cognitive functioning.

In contrast, kinesthetic measures of movement remained largely unaffected by load – reach duration and trajectory length did not change with load, and reach speed showed only a weak omnibus effect without significant pairwise differences. Given evidence that WM maintenance and motor planning may rely on shared neural resources [20–26], we hypothesized that increasing load would consume more of this shared capacity, thereby impairing motor preparation and subsequent execution. Consistent with this view, higher load prolonged motor preparation, as reflected by longer pre-reach durations, yet movement execution itself remained intact. These findings indicate that increasing load primarily burdens cognitive and planning stages before movement onset, distinguishing cognitive preparation from motor execution dynamics [57]. In other words, memory load exerts its strongest influence prior to movement, where once the movement is initiated, performance is relatively protected from additional WM loads.

### 4.2. Parieto-Occipital Activity Scales with Load and Exhibits Posterior-Frontal Engagement

We confirmed that posterior activity scaled robustly with load during both the early visual-evoked N1 and the following delay period. At parieto-occipital sites, greater negativity accompanied higher loads, while frontal and central sites showed complementary positivity. This engagement aligns with models positing that the posterior sensory-perceptual cortical regions support maintenance functions of WM, whereas PFC regions provide top-down control, including activation of goals, inhibition of distractors, and response preparation [58–60]. Although our study does not localize specific cortical generators, the observed posterior-frontal engagement is consistent with the frontoparietal network engagement in visual processing, WM maintenance, and subsequent action planning [61–63]. Our topographical scalp maps further illustrate that relative to the baseline 1-disc condition, early visual interval (N1) showed primarily frontal engagement at a moderate load (3-disc – 1-disc), followed by additional posterior involvement at a higher load (4-disc – 1-disc). During the delay interval, the pattern reversed – posterior engagement emerged at a modest load (e.g., 2-disc – 1-disc), and additional frontal recruitment appeared at higher loads (3- and 4-disc – 1-disc), consistent with prior reports of greater frontal contribution under higher demands [15–19].

Further, contralateral signals carried the strongest load effects. During the N1 interval, activity increased linearly with load, consistent with enhanced visual attention and perceptual processing as the number of visual items increased [46, 64]. During the delay period, both contralateral activity and the CDA (i.e., contralateral – ipsilateral) scaled with load but reached an asymptote at 3-discs, particularly when averaged across posterior electrodes. This replicates classic visual WM capacity signatures [3, 9, 47, 65]. Importantly, this plateau diverged from behavior, such as ED, which continued to worsen beyond 3-discs. Prior work similarly identified PPC regions, especially in the intra-parietal and intra-occipital sulci, as reflecting visual WM storage capacity, where activity saturated at capacity even as behavioral accuracy declined and reaction times lengthened [13]. Together, these results demonstrate that posterior contralateral activity in our task reliably indexes mnemonic load prior to movement, with only delay period activity reaching an asymptote at capacity and reflecting memory limits rather than task difficulty.

### 4.3. Posterior Load Effects Predict Behavioral Performance

We found that CDA amplitude related to behavioral performance only at a moderate memory demand (2-disc), likely near the task-related capacity limit. At this load, greater CDA negativity during the delay period correlated with better WM performance (smaller ED) but slower reach speed, suggesting a potential tradeoff between cognitive and motor performances indexed by the same neural signature. This selective brain-behavior coupling indicates that CDA is most informative when performance is neither at ceiling nor at floor. At higher loads, neural signals likely saturate and the increased reliance on guessing introduces behavioral variability that obscures correlations, and at lower loads, mnemonic demands are too minimal to produce meaningful variance. Standard neuropsychological measures converged with these findings, where higher Digit Span scores correlated with better WM performance (smaller ED) again at moderate loads (2- and 3-disc). Together, these converging behavioral and neuropsychological findings highlight posterior delay period activity as a useful task-related neural marker of performance, particularly at moderate mnemonic loads near capacity.

### 4.4. Handedness Exerts Selective Behavioral but Not Neural Effects

Using the non-dominant hand may impose additional planning and control demands [66, 67] prior to movement, particularly under higher WM load. We therefore hypothesized that in our action-coupled WM task, such asymmetries might manifest in both motor preparation and behavioral performance, as well as in underlying neural activity. Behaviorally, response with the dominant hand resulted in faster reach speeds but prolonged pre-reach durations, suggesting more deliberate preparation followed by better control during execution. According to the complementary dominance model of lateralization, where the dominant and non-dominant hands are proposed to be specialized for different aspects of motor control, the dominant hand outperforms the non-dominant hand with increased task repetition, demonstrating efficient adaptation, whereas the non-dominant hand may prevail when task experience is at the initial phase [68]. Therefore, future trial-by-trial investigations could provide further evidence for this model. Past work has also shown that the dominant hand excels in precision and dynamic control, whereas the non-dominant hand exhibits greater movement variability [67, 69]. Consistent with this, we observed greater response dispersion when the non-dominant hand reached contralaterally, compared to the dominant hand. Despite these behavioral asymmetries, posterior neural signals tracking load were unaffected by handedness, suggesting that these pre-movement neural signals are invariant to motor demands. This implies that both hands recruit comparable neural resources to meet task demands and that our paradigm may not vary motor demands enough to elicit dominance-related differences in neurotypical adults. Therefore, future work should test whether this invariance holds under broader variations in motor demands alongside differing cognitive loads.

### 4.5. Summary and Future Directions

In this study, we showed that increasing load impaired WM performance and prolonged the preparatory phase of action while sparing motor execution; task performance engaged frontoparietal networks, with posterior delay activity saturating at capacity; and load-related neural activity most strongly linked to behavior at moderate loads near capacity. Handedness selectively influenced behavior but did not modulate posterior load-related neural activities. Together, these findings provide insight into how mnemonic demands shape the neural basis of cognitive-motor interactions prior to behavioral responses and identify where such neural signatures most effectively index behavior. Our handedness analysis included only right-hand dominant participants, but in future work, it will be interesting to also include left-hand dominant participants to potentially strengthen generalizability and deepen understanding of the neural basis of handedness, considering both unique and shared hemisphere-specific activities, in the scope of WM-guided actions. In addition, incorporating additional markers (e.g., index finger) and extending analyses to the peri-movement phase in future work could provide a more comprehensive picture of upper extremity kinematics and the neural bases of task-related cognitive-motor integration. Lastly, beyond the temporal precision of EEG, future work combining this paradigm with source localization or task-based fMRI techniques could also help better delineate the spatial origins and spatiotemporal dynamics of action-coupled load effects. Overall, our study, which couples visual WM demands with dynamic upper extremity responses in a single task, provides a translational framework for investigating cognitive-motor interactions and establishes a foundation for potentially examining how these processes may be disrupted in neurological conditions characterized by both cognitive and motor impairments, such as Parkinson’s disease.

## DATA SHARING STATEMENT

Data and code from this study will be available through a public repository upon publication. The authors will work with the editorial office during production to incorporate appropriate links.

## CREDIT AUTHORSHIP CONTRIBUTION STATEMENT

**Jeehyun Kim**: Conceptualization, Data Curation, Formal Analysis, Investigation, Methodology, Writing – Original Draft. **Edward G. Freedman**: Conceptualization, Funding Acquisition, Methodology, Project Administration, Supervision, Writing – Original Draft. **John J. Foxe**: Conceptualization, Funding Acquisition, Methodology, Project Administration, Supervision, Writing – Original Draft.

## ACKNOWLEDGMENTS

We would like to thank each of the participants that enrolled in the study. Special thanks to Dalia Einstein, Matthew Cowie, Aseem Dani, Emma Mantel, and Dr. Gil Rivlis for their help with this study.

## ABBREVIATIONS LIST

3D: three-dimensional
ANOVA: analysis of variance
CDA: contralateral delay activity
ED: error distance
EEG: electroencephalography
ERP: event-related potential
ICA: independent component analysis
LSL: Lab Streaming Layer
LxL: left target hemifield-left response-hand
LxR: left target hemifield-right response-hand
MAD: median absolute deviation
MoBI: mobile brain-body imaging
MoCA: Montreal Cognitive Assessment
PFC: prefrontal cortex
PPC: posterior parietal cortex
rCV: robust coefficient of variation
RxL: right target hemifield-left response-hand
RxR: right target hemifield-right response-hand
SCP: statistical cluster plot
SD: standard deviation
WAIS: Wechsler Adult Intelligence Scale
WM: working memory

## Clinical Trial Number

Not applicable.

## Consent for publication

Not applicable.

## Ethics approval and consent to participate

All aspects of the research conformed to the tenets outlined in the Declaration of Helsinki, with the exception that this study was not preregistered. The institutional review board of the University of Rochester, where the data collection took place, approved this study (STUDY00001952). All participants provided written informed consent.

---

**Supplementary Figure 1.**
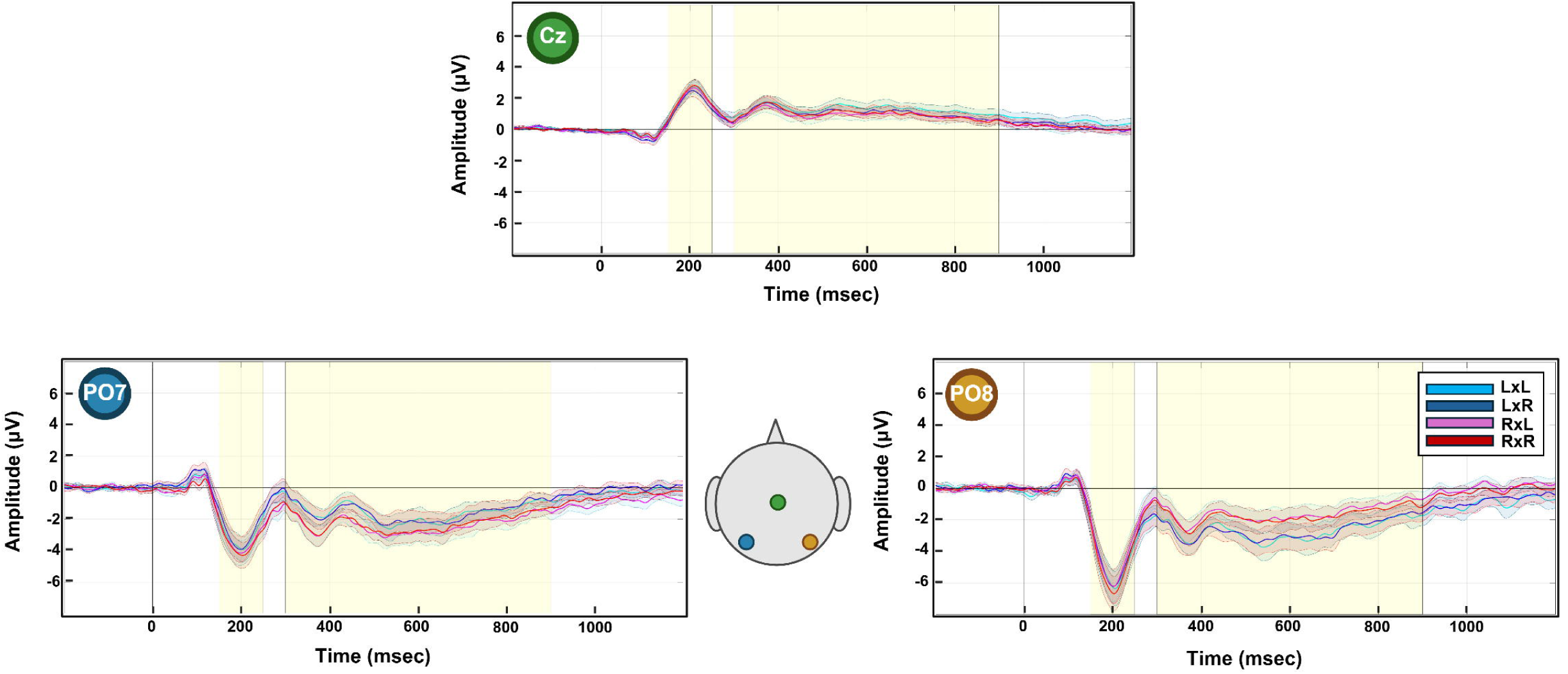
Neurophysiological Activities by Target Hemifield and Response-Hand. The average waveforms with SD in colored shades are time-locked to the encoding array onset (time 0) across different conditions (i.e., target hemifield and response-hand pairs). Highlighted regions correspond to the intervals selected for preplanned statistical analysis. Colored circles indicate scalp location for each corresponding waveform.

**Supplementary Figure 2.**
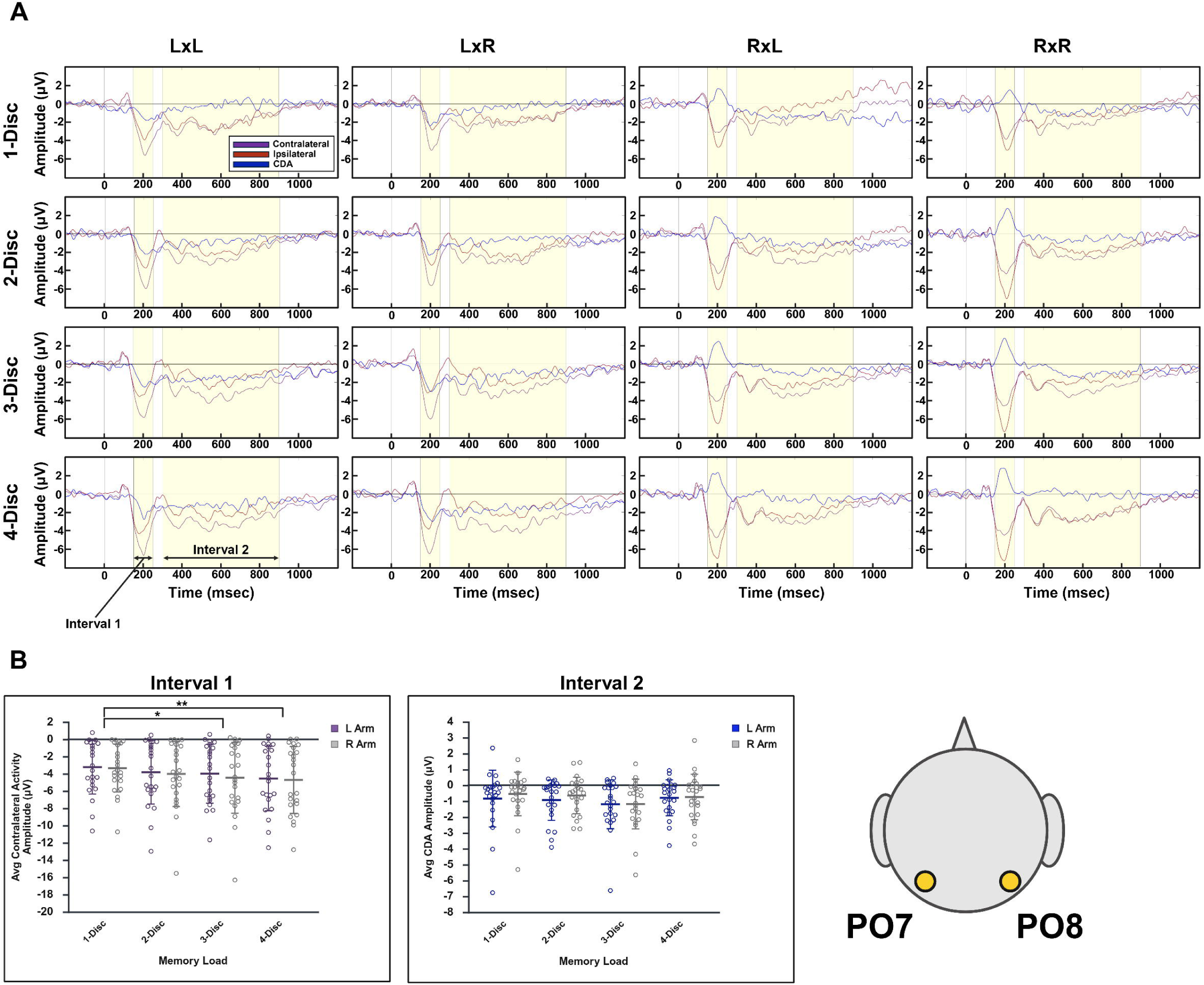
Neurophysiological Activities by Different Conditions Across Loads Using PO7/PO8 Pair. (A) shows average contralateral (purple), ipsilateral (red), and CDA (i.e., contralateral minus ipsilateral difference; blue) waveforms time-locked to the encoding array onset (time 0) across different memory loads for each condition (i.e., target hemifield and response-hand pair). Highlighted regions correspond to the intervals selected for preplanned statistical analysis. For LxL and LxR conditions, we probed contralateral activity from PO8, and for RxL and RxR conditions, we probed contralateral activity from PO7. In (B), unfilled circles show individual participant means for contralateral activity (purple: left response-hand; grey: right response-hand) during interval 1 (left panel) and for CDA (blue: left response-hand; grey: right response-hand) during interval 2 (right panel). The central line and error bars indicate group means +/- SD. * indicates *p* < 0.05. ** indicates *p* < 0.01.

